# SIP-BERT: A multi-organism deep strategy for predicting self interaction in proteins

**DOI:** 10.64898/2026.02.04.702782

**Authors:** Tapas Chakraborty, Saikat Majumder, Padmalochan Maiti, S.V.S.S.N.V.G. Krishna Murthy, Anup Kumar Halder, Subhadip Basu

## Abstract

Self-interacting proteins (SIPs) are critical to cellular regulation, yet their experimental identification remains challenging due to high costs, inefficiencies, and frequent false positives. Leveraging recent advances in deep language models, we introduce SIP-BERT, a family of lightweight transformer-based models trained on organism-specific self-interaction datasets curated from existing protein–protein interaction databases. We developed three variants: SIP-BERT(H), SIP-BERT(Y), and SIP-BERT(HY) -trained on human, yeast, and combined datasets, respectively. These models significantly outperform existing methods, exceeding baseline accuracies by 18%, 8% and 15% respectively. SIP-BERT models also generalise effectively to under-annotated organisms such as the mouse and the fruit fly, achieving high recall despite minimal labeled data. Furthermore, structural analysis of predicted false positives using PDB-derived alpha-carbon distance maps reveals close spatial residue proximities, suggesting plausible but undocumented self-interactions. These results highlight the potential of SIP-BERT to uncover novel SIPs and expand our understanding of protein self-interaction across diverse species. The dataset and the developed models are available at https://github.com/CMATERJU-BIOINFO/SIP-BERT for academic use only.

## 1 Introduction

Proteins are fundamental biological macromolecules present in all living organisms. They play a crucial role in numerous cellular processes [1], including metabolism, signal transduction, hormone regulation, DNA transcription, and replication [2]. Most proteins carry out their functions by interacting with other proteins, making protein-protein interactions (PPIs) essential to understanding cellular activity. Also, identification and analysis of PPI data offer valuable insights into the structural characteristics [3] of proteins and the complex molecular machinery operating within cells [4].

A significant subset of protein-protein interactions (PPIs) involves self-interacting proteins (SIPs)-proteins that bind to identical copies of themselves, typically encoded by the same gene. These self-interactions often result in the formation of homo-oligomers, which are essential for various biological functions, including enzyme activation, signal transduction, immune responses, and regulation of gene expression [5].

Comprehending how proteins interact is vital to deciphering cellular behaviour and molecular mechanisms. Computational approaches usually simplify these interactions as binary relationships between protein pairs. Organisms, such as yeast and humans, have well-established networks of experimentally validated protein-protein interactions (PPIs). Insights derived from these networks have played a pivotal role in advancing our understanding of functional genomics [6] and enabling in-depth analysis of biological pathways [7].

### 1.1 Existing Works on PPI and SIP

*In-vivo* experimental methods for detecting PPIs have been used for years [8], [9], [10], [11], [12], [13], [14] which are time-consuming and error-prone. On the other hand, computational methods leveraging artificial intelligence and machine learning became popular due to the availability of large volume of protein sequence information from advanced technologies. Protein sequences are crucial for *in-silico* PPI prediction methods, and sequence encoding is the key attribute of these approaches.

Zahiri et al. [15] proposed PPIevo, an amino acid sequence-based model that uses a position-specific scoring matrix (PSSM) to extract evolutionary information, leading to improved prediction accuracy. Many researchers have used protein sequences as the basis of PPI prediction [16], [17], [18]. Some of them used traditional models like SVM, some researchers prefer using deep neural networks, combined with projection and data augmentations, to predict interactions. DensePPI [19] was published a couple of years ago and uses a unique encoding strategy. They have used a random colour map to generate images of the two interacting or non-interacting proteins. These rectangular images were used to train a deep learning model (DenseNet201) to for protein interaction prediction.

DL-PPI [20], another PPI method, also uses sequence data for predicting protein–protein interactions (PPIs). It incorporates two key modules designed to enhance feature extraction from individual protein sequences and to capture interactions among proteins in unfamiliar datasets. First, the Protein Node Feature Extraction Module employs the Inception method to efficiently extract meaningful patterns and representations from protein sequences, improving feature quality. Second, the Feature-Relational Reasoning Network (FRN) utilizes Graph Neural Networks within the Global Feature Extraction module to analyze interactions between protein pairs, capturing complex relational information.

Recent research, reflected across several key publications, demonstrates significant progress in both the biological understanding of SIPs and the computational methods developed for their prediction. Building on this understanding, several computational approaches have been developed to predict SIPs from primary sequence data. Techniques range from recurrent neural networks (RNNs) leveraging evolutionary features to ensemble learning models (e.g., PSPEL), which integrate diverse classifiers for robust predictions. The use of random projection and Fast Fourier Transform introduces novel signal processing techniques to extract interaction-relevant features from sequences [21], [22], [23], [24], [25]. Further refinement is evident in methods such as SPAR [26], which applies random forest classifiers enriched with fine-grained domain annotations for enhanced accuracy and scalability. Together, these works highlight a trend towards integrative, machine learning-based models that combine sequence, structural, and evolutionary data. They represent a paradigm shift from traditional experimental approaches to in silico prediction.

Modern deep neural network architectures—particularly those designed for sequence data, like BERT [27] combined with large-scale pre-training, have transformed the landscape of automated text analysis. Among them, the Transformer architecture, built around attention mechanisms, has set new performance standards across a wide range of benchmarks and application domains. Motivated by BERT’s success in natural language processing (NLP), researchers have recently adapted it for biological sequence analysis [28], [29]. Protein-BERT is a domain-specific variant of the BERT architecture, optimized for protein sequence modelling. It harnesses BERT’s powerful pre-training and transfer learning capabilities to capture the complex biochemical and structural properties of proteins. Like its NLP counterpart, Protein-BERT consists of multiple layers of self-attention and feed-forward neural networks, but with a tailored encoding mechanism designed to recognize long-range dependencies and evolutionary patterns in amino acid sequences. Its bidirectional nature allows the model to consider both upstream and downstream amino acids when encoding a given position, enabling a nuanced understanding of protein structure and interactions.

Despite significant progress, current approaches still face notable limitations. Most proteinprotein interaction (PPI) prediction models are not well-suited for identifying self-interacting proteins (SIPs), and their performance in this context is often suboptimal. Moreover, the development of computational models specifically targeting SIPs remains limited, with many existing methods overlooking key challenges such as imbalanced datasets. Addressing these issues is crucial for enhancing prediction accuracy. Therefore, there is a pressing need to explore more efficient and robust computational strategies tailored to SIP prediction. High-quality negative data, i.e. non-self-interaction data, is essential for model training. Unavailability of such data makes this research more challenging.

### 1.2 Present Work

Comprehensive datasets specific to self-interacting proteins (SIPs) are limited in availability, and existing studies often rely on earlier versions of protein–protein interaction (PPI) resources. To facilitate robust model development and evaluation, we propose curating a benchmark dataset by integrating the most recent versions of widely used PPI databases, including BioGRID [30], DIP [31], STRING [32], and MINT [33].

In the current study, we build upon the curated dataset to introduce SIP-BERT (Self-Interaction Protein BERT), a deep language model specifically designed for predicting self-interacting proteins (SIPs). To address both species-specific and cross-species SIP prediction, we propose the development of three tailored variants of SIP-BERT: SIP-BERT(H), trained exclusively on human self-interaction data; SIP-BERT(Y), trained on yeast-specific data; and SIP-BERT(HY), a hybrid model trained on a combined human-yeast dataset to improve cross-species generalizability. These models will be further evaluated on underrepresented organisms such as Drosophila melanogaster (FruitFly) and Mus musculus (Mouse), where limited labeled data restricts conventional training. Through this approach, we aim to demonstrate the robustness, scalability, and evolutionary transferability of the proposed SIP-BERT framework.

## 2 Methodology

This study presents a deep learning framework for predicting Self-Interacting Proteins (SIPs) by leveraging a BERT-based architecture to jointly represent protein sequences and Gene Ontology (GO) annotations. The methodology section is structured into four main components: Benchmark Data Preparation, Preprocessing & Input Embedding, Deep Learning Model Architecture, and the Classification Layer with Strategy. An overview of the complete workflow is provided in Figure 1, and each stage is elaborated in the subsequent sections.

**Fig. 1.**
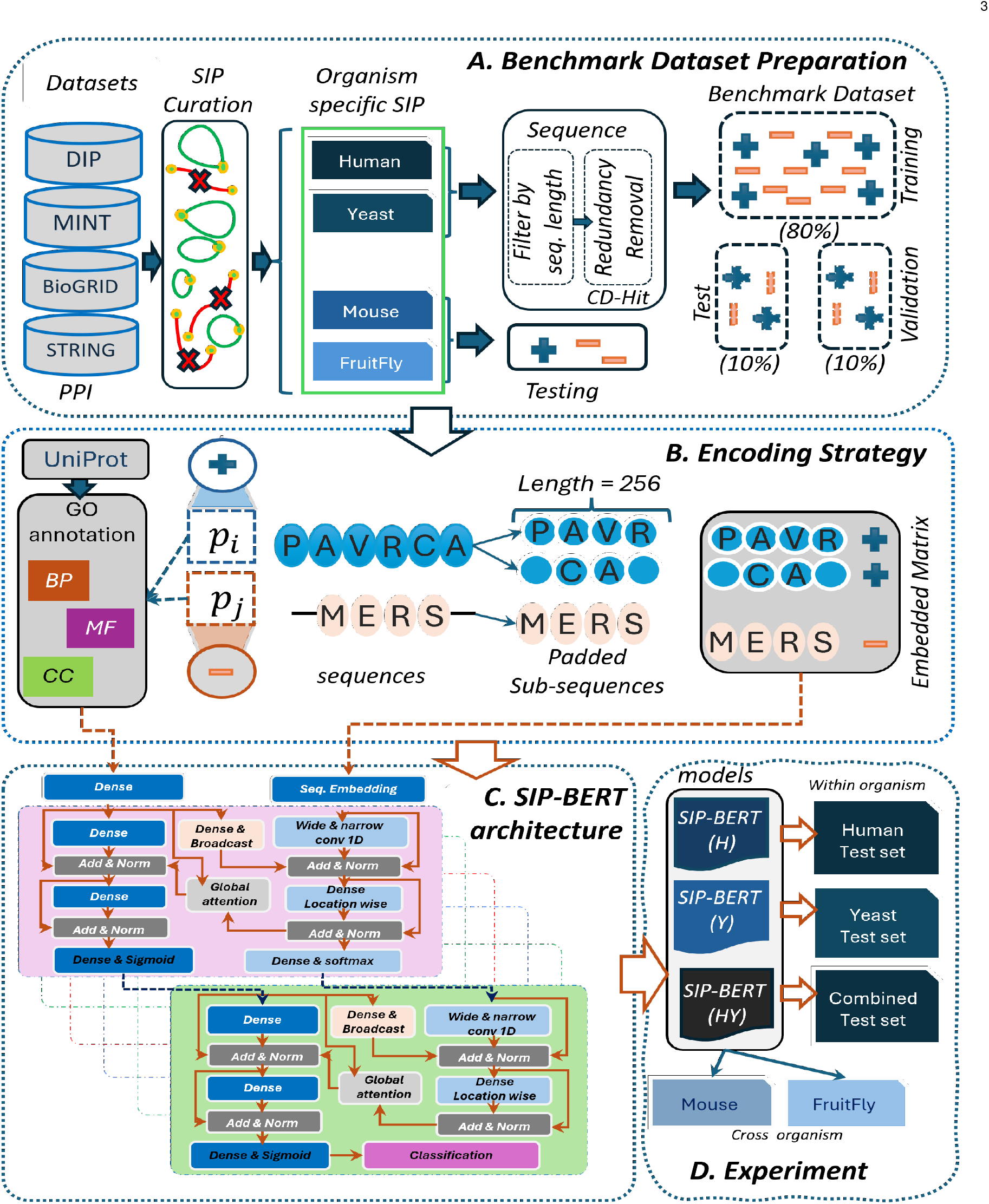
SIP-BERT workflow: A pictorial representation of the protein self-interaction prediction using SIP-BERT. A) Benchmark dataset preparation: Self-interacting proteins of human, Yeast, and other organisms have been taken from the latest version of PPI datasets. Sequences of length between *128* and *5000* are considered. Redundant sequences (similarity *≥*80%) are discarded using CD-hit [34]. B) Encoding strategy: Sequences of length more than 256 have been divided into multiple sequences. Shorter sequences have been padded with <PAD> tokens on either side. Few additional tokens (*<*START*>, <*END*>*, etc) have been used for encoding. C) Model architecture: Input sequence and GO annotation (biological process, molecular function, cellular component) used to train the BERT model. The architecture partitions the network into two almost parallel blocks to facilitate global and local representations. D) Experiment: BERT model was trained on benchmark datasets to develop three new models: SIP-BERT(H), SIP-BERT(Y) and SIP-BERT(HY). Performance of these models was tested on the held-out set. Cross-organism performance was also evaluated on organisms where the training data is too small, e.g. for mouse and fruitfly.

### 2.1 Benchmark Data Preparation

Self-interaction data have been taken from the latest version of BioGRID [30], DIP [31], STRING [32], and MINT [33]. Organism-wise count of self-interacting and non-self-interacting proteins given in Table 1

**TABLE 1.** Organism-wise count of self-interacting and non-self-interacting proteins.

| Organisms | SIP | nonSIP |
| --- | --- | --- |
| Human | 3617 | 12797 |
| Yeast | 1836 | 2220 |
| Mouse | 682 | 9398 |
| FruitFly | 360 | 2251 |

In this experiment, protein sequences and corresponding Gene Ontology (GO) annotations for the curated dataset are obtained from the UniProt database [35]. As part of our pre-processing pipeline, we selected protein sequences ranging from *128* to *5000* amino acids in length. To ensure dataset diversity and minimize redundancy, sequences with *≥*80% identity were filtered out using CD-HIT [34]. Each sequence is represented as non-overlapping segments of fixed length 256. Sequences shorter than 256 residues are symmetrically padded with ⟨PAD⟩ tokens to maintain uniform input size. This standardization ensures consistent input representation for downstream modeling.

To construct a balanced training dataset for model development, a curated set of protein sequences from Saccharomyces cerevisiae and Homo sapiens was used. In the case of yeast, a total of 6,622 unique proteins were obtained from the UniProt database, among which the number of self-interacting proteins (SIPs) was substantially lower than that of non-self-interacting proteins (non-SIPs). Interestingly, SIPs exhibited a higher average sequence length compared to non-SIPs. To address the inherent class imbalance, non-SIP instances were randomly downsampled to match the number of SIPs, ensuring an equal representation of both classes during training. The resulting yeast dataset comprised 1,836 SIPs and 2,220 non-SIPs. A similar curation and balancing strategy was employed for the human dataset, derived from 16,414 unique proteins, leading to a final benchmark set of 3,288 SIPs and 4,847 non-SIPs. These datasets were later combined to form a comprehensive benchmark comprising 5,088 SIPs and 8,847 non-SIPs. An 80:10:10 split was employed for training, validation, and testing, respectively, to support effective model development and unbiased performance assessment

### 2.2 Preprocessing & Input Embedding

#### Protein sequence tokenization

Each protein sequence is represented as a linear string composed of standard amino acid residues. Let

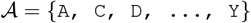

denote the set of the 20 canonical amino acids. To accommodate sequence boundary definitions, padding, and non-standard or unknown residues, we augment this set with special tokens: ⟨START⟩, ⟨END⟩, ⟨ PAD⟩, and ⟨OTHER⟩.

Each protein sequence is subsequently tokenized into a fixed integer-based vocabulary, resulting in the following structured representation:

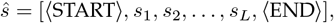

where *s*_*i*_ *∈A* for 1 *≤ i ≤ L*, and *L* denotes the sequence length.

To accommodate fixed-length batch processing, we pad each sequence symmetrically if its length is less than a fixed maximum length *N*_*max*_ = 254:

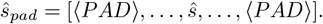

By padding equally on both sides of the sequence, the subsequence stays centered, allowing convolutions to effectively capture positional context for shorter sequences.

#### GO-annotation encoding

Gene Ontology (GO) annotations effectively consolidate the counts of three types of gene ontology annotations into a single array for each protein, which is then used as input to the model. This allows the model to leverage additional biological information alongside the sequence data, enhancing its predictive capabilities.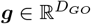 denote this vector.

### 2.3. Deep Learning Model

The proposed architecture consists of 6 encoders(*N*_*enc*_ = 6) stacked sequentially to enable hierarchical and deep feature extraction prior to classification. Every encoder block integrates the sequence features and GO annotation simultaneously by processing the embedded protein sequence along with its Gene Ontology (GO) annotation representation(*h*_*global*_). The sequence branch extracts local and long-range dependencies by employing parallel one-dimensional convolutions: a narrow convolution for local dependencies and a wide convolution for detection of long-term connection. The outputs of these convolutions are then projected through dense layers and concatenated with the GO-based representation(*h*_*global*_) through residual connections. This concatenated representation is then normalized and further refined using a position-wise feedforward dense layer with dropout followed by an additional residual normalization. Subsequently, a global attention mechanism allows the global representation(*h*_*global*_) to selectively pay attention to the sequence tokens, and the output of the produced attention is concatenated back to the global representation(*h*_*global*_) through additional residual transformations. Each of the first five encoder blocks generates two outputs: a sequence-level prediction from a dense layer with softmax activation applied to the refined sequence embedding, and an output annotation(*h*_*global*_) prediction from a dense layer with sigmoid activation applied to the updated global representation(*h*_*global*_). The final (sixth) encoder block outputs only the classification score based on the output annotation.

#### 2.3.1. Dense Layer for GO annotation

In parallel with the sequence embedding pathway, the model processes the Gene Ontology (GO) annotation vector of each protein to generate a dense global representation. This global embedding encapsulates high-level functional and contextual information, which is subsequently integrated with local sequence-derived features through feature fusion and attention mechanisms. The fusion facilitates the model’s capacity to jointly capture biological semantics at both global and local scales.

Let the fixed-length GO annotation vector for a protein be denoted as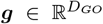 Prior to entering the encoder blocks, this vector is projected into a rich, high-dimensional feature space:

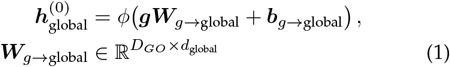

where *d*_*global*_ is the dimensionality of the global representation (e.g. *d*_*global*_ = 512), and *ϕ* denotes the nonlinear activation (e.g. GELU). A dropout layer may optionally follow this projection:

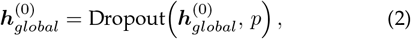

where *p* denotes the dropout rate.

This densely transformed annotation vector 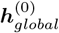 serves as the hidden state of the global branch in the input stage. Throughout each encoder block, this global vector is updated after receiving information from the sequence branch via attention and feature fusion.

#### 2.3.2 Convolutions for Local and Global dependencies

In protein sequence consecutive amino acids (motif) may contribute significantly to that protein characteristics and functionality [29]. Inspired by the analogy between letters, words, and sentences, we recognize that single amino acids lack meaning in isolation, but blocks of consecutive residues can capture informative patterns. To extract such motifs at different scales, we feed the embedded sequence ***E***_*seq*_ into two parallel one-dimensional convolutional paths: i)**Narrow Convolution:** A 1D convolutional layer with kernel size *k*_*narrow*_ = 9, dilation rate 1, and stride 1 captures local subsequence features similar to ‘words’ in natural language. Proteins do not have explicit ‘words’, so a kernel of 9 allows the model to look at 9 consecutive amino acids and captures local patterns in the sequence. ii) **Wide Convolution:** Another 1D convolutional layer with the same kernel size (*k*_*wide*_ = 9) but a dilation rate of 5 captures long-range dependencies across the sequence:

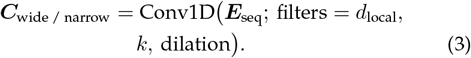

##### Dropout

Dropout layers are applied after each convolution to prevent overfitting by randomly setting a fraction of the input units to zero during training. Both ***C***_*narrow*_ and ***C***_*wide*_ pass through Dense and Dropout layers:

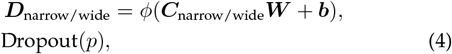

where 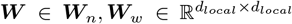 and *p* denotes the dropout rate. This mapping ensures both paths output features in a shared representation space.

#### 2.3.3 Residual Connections and Layer Normalization

##### Feature Fusion

The global representation ***h***_*global*_ into the sequence branch via a Dense projection, narrow convolution, and wide convolution are combined with the original sequence representation using a feature fusion. This residual connection helps in training deeper networks by allowing gradients to flow more easily.

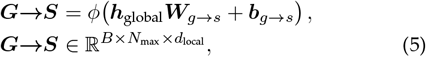

##### Layer Normalization

After the fusion, layer normalization is applied to stabilize the learning process and improve convergence.

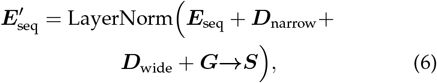

followed by a position-wise Dense layer with Dropout:

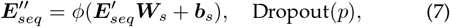

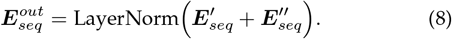

#### 2.3.4 Global Attention Mechanism

The **Global Attention** mechanism enhances the model’s ability to integrate sequence-level features with global contextual information. It aligns each token in the sequence with a fixed global representation, enabling selective focus on relevant regions of the sequence conditioned on the overall context.

##### Inputs

###### Global Representation (*X*)

A fixed-dimensional vector summarizing contextual information, derived from auxiliary encoders or pooled features.

###### Sequence Representation (*S*)

A matrix of token-level embeddings, with each row corresponding to a sequence element (e.g., amino acid).

##### Attention Computation

###### Query (*Q*)

Computed from the global vector using weight matrix *W*_*q*_, producing a tensor of shape (batch_size, *n*_heads_, *d*_key_). It represents the model’s query with respect to the global context.

###### Keys (*K*)

Generated from the sequence embeddings using *W*_*k*_, resulting in shape (batch_size, *n*_heads_, seq_len, *d*_key_). Keys describe each sequence position in terms of its relevance to the query.

###### Values (*V*)

Also derived from sequence embeddings using *W*_*v*_, with the same shape as the keys but with dimensionality *d*_value_. Values carry the sequence features that will be aggregated.

##### Attention Weights

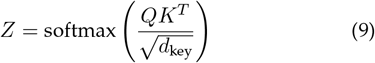

This produces weights for each sequence position relative to the global query.

##### Contextual Output

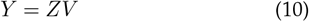

A weighted sum of values across all sequence positions. The outputs from all attention heads are concatenated into a tensor

###### Fusion into the Global Branch

The attention output ***A*** is combined with the previous global state via a multi-step residual update for an enhanced context-aware global representation:

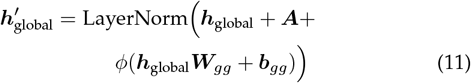

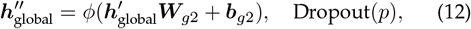

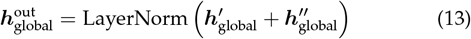

This residual fusion integrates the sequence-aware attention output into the global feature vector while preserving high-level contextual integrity.

By stacking six blocks, the receptive field of the wide convolution reaches 241 residues. This enables the model to capture both the global annotation context and local sequence patterns for rich, context-aware representations.

### 2.4 Classification Layer

#### Intermediate Outputs

For each encoder block (except the last), the model produces output sequence (using a dense layer with softmax activation) and for output annotations (using a dense layer with sigmoid activation).

#### Final Output Layer

To predict a binary classification from the last encoder block, we take the global representation 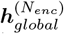 (output annotation) and pass it through a final Dense layer with sigmoid activation. This output can be used for tasks such as classifications or regression.

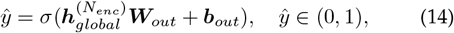

which predicts the probability a protein is self-interaction (Binary Classification).

### 2.5 Classification Strategy

To evaluate the performance of the proposed model against *state-of-the-art* approaches, we employ a held-out test set that is not used during training or validation. As the prediction is performed at the sub-sequence level, we adopt two aggregation strategies to derive the final prediction at the full protein sequence level: the *Average Strategy* and the *Consensus Strategy*.

#### Average Strategy

The test data is evaluated using ten independently trained models, and the mean prediction score is computed across these models. The final classification is determined based on the averaged output probabilities.

#### Consensus Strategy

Here, 1-hot rule is used. Held out set given to ten models to get five classification results. Final prediction result is generated using 1-hot rule *FC*_*k*_.

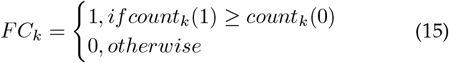

## 3 Results and Discussions

The effectiveness of all the SIP-BERT models is assessed using a held-out set of the following benchmark datasets: a) *S. cerevisiae* dataset and b) Human dataset c) combination of *S. cerevisiae* and Human dataset. SIP-BERT model’s learning is configured as per the parameters described in Figure 2.

**Fig. 2.**
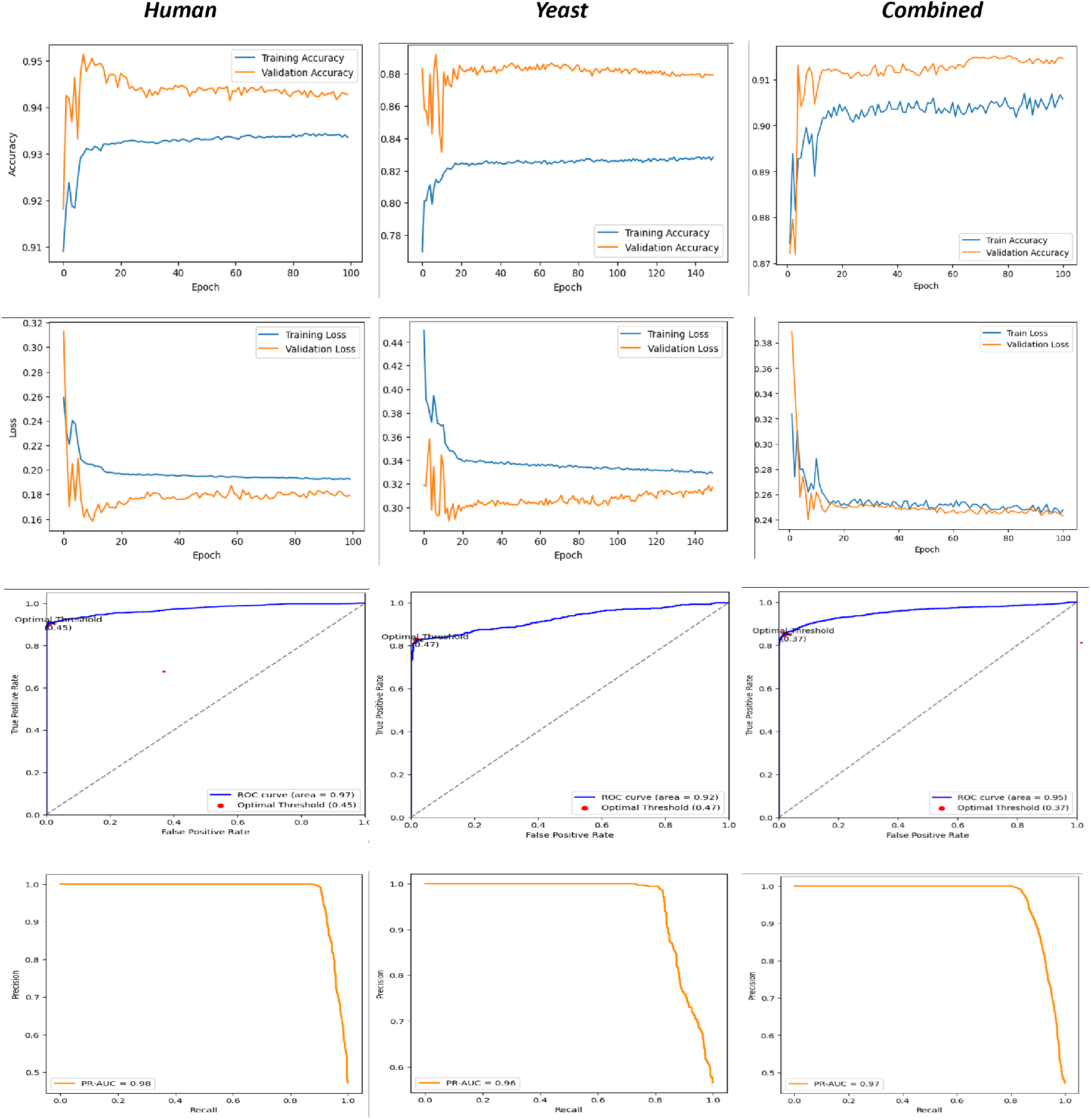
Performance evaluation of three models: *SIP-BERT(H),SIP-BERT(Y) and SIP-BERT(HY)* A pictorial representation of the performance of the organism-specific SIP-BERT models. A) First set of graphs represent training and validation accuracies over epochs. B) The second set of graphs represents training and validation loss over epochs. C) The third set of graphs represents True positive vs false positive rates. D) The last graphs represent the precision-recall curve.

Model evaluation is conducted using performance metrics-Area under the Receiver Operating Curve (AUC), Precision (Prec), Recall, F-score and Accuracy (Acc). Comparative results, showcasing the model’s alignment with *state-of-the-art* methods, are presented in Table 2, 3 and 4 for all three datasets, respectively.

**TABLE 2.** Performance comparison of SIP-BERT(Y) with baseline methods on the *S*.*cerevisiae* dataset.

| Methods | Acc | Prec | Recall | F1-score | AUC |
| --- | --- | --- | --- | --- | --- |
| DensePPI [19] | 55.00 | 57.00 | 56.00 | 53.00 | 56.00 |
| DL-PPI [20] | 52.00 | 53.10 | 55.30 | 54.20 | - |
| Li et al [5] | 65.02 | 65.00 | 65.00 | 65.00 | 65.00 |
| ProteinBERT [29] | 73.08 | 68.52 | 70.59 | 69.06 | 79.21 |
| SIP-BERT(Y) | 81.41 | 85.47 | 68.20 | 76.20 | 85.64 |

**TABLE 3.** Comparative performance of SIP-BERT(H) and existing methods on the human self-interaction prediction task.

| Methods | Acc | Prec | Recall | F1-score | AUC |
| --- | --- | --- | --- | --- | --- |
| DensePPI [19] | 58.00 | 58.00 | 58.00 | 58.00 | 58.00 |
| DL-PPI [20] | 60.02 | 60.73 | 93.20 | 73.56 | - |
| Li et al [5] | 61.50 | 60.00 | 62.00 | 58.00 | 66.00 |
| ProteinBERT [29] | 72.65 | 64.91 | 70.37 | 67.48 | 77.70 |
| SIP-BERT(H) | 90.81 | 96.27 | 81.22 | 88.08 | 92.99 |

**TABLE 4.** Comparative evaluation of SIP-BERT(HY) and existing methods on a unified benchmark dataset combining human and *S. cerevisiae* proteins.

| Methods | Acc | Prec | Recall | F1-score | AUC |
| --- | --- | --- | --- | --- | --- |
| DensePPI [19] | 60.00 | 58.00 | 60.00 | 58.00 | 60.00 |
| DL-PPI [20] | 56.99 | 58.34 | 87.02 | 69.85 | - |
| Li et al [5] | 65.00 | 61.00 | 61.00 | 61.00 | 65.00 |
| ProteinBERT [29] | 71.08 | 65.70 | 67.74 | 66.52 | 76.62 |
| SIP-BERT(HY) | 86.34 | 91.53 | 75.39 | 82.43 | 90.12 |

### 3.1 Performance on *S*.*cerevisiae* Dataset

We evaluated the performance of SIP-BERT(Y) on the *S. cerevisiae* dataset and compared it against a range of established methods, including traditional PPI prediction models and SIP-specific approaches. As shown in Table 2, generic and recent PPI models such as DensePPI and DL-PPI yielded modest performance, with accuracies of 55.00% and 52.00%, respectively, and F1-scores below 55%, indicating limited effectiveness in identifying self-interacting proteins (SIPs). The SIP-focused method proposed by Li *et al*. [5] provided a marginal improvement, achieving 65.02% accuracy and an AUC of 65.00%, but still struggled to generalize effectively.

To further contextualize performance, we compared SIP-BERT(Y) with ProteinBERT [29], a recent protein language model incorporating evolutionary context. ProteinBERT achieved the highest performance among existing baselines, with 73.08% accuracy and an AUC of 79.21%. However, SIP-BERT(Y) consistently outperformed all benchmark models, achieving 81.41% accuracy, 76.20% F1-score, and an AUC of 85.64%. This improvement suggests that SIP-BERT(Y) effectively captures sequence-level patterns specific to SIPs, offering a more tailored and discriminative representation compared to general-purpose models.

### 3.2 Performance on Human Dataset

Evaluation on the human benchmark dataset reinforces the effectiveness of SIP-BERT(H). As shown in Table 3, the model achieved 90.81% accuracy and an AUC of 92.99, markedly outperforming all baseline methods. Specifically, it achieved an 18% absolute improvement in accuracy over ProteinBERT (72.65%), the best-performing existing approach. Additionally, SIP-BERT(H) attained higher recall (81.22%) and F1-score (88.08%), reflecting its strong discriminative ability in identifying human self-interacting proteins. Collectively, the consistent gains observed across all evaluation metrics in Table 3 underscore the robustness and superior predictive capacity of SIP-BERT(H) compared to both conventional and contemporary deep learning-based methods.

### 3.3 Performance on Combined Dataset

Table 4 presents model performance on the combined dataset of S. cerevisiae and human proteins. SIP-BERT(HY) achieved an accuracy of 86.34%, improving over Protein-BERT by approximately 15%, with corresponding gains across precision, recall, F1-score, and AUC.

Despite the increased data volume, SIP-BERT(HY) performed slightly below its human-specific counterpart, SIP-BERT(H). Compared to SIP-BERT(H), accuracy decreased by 4%, F1 score by 6%, and AUC by 2%, suggesting that training of the cross-organism introduces additional variability, which can affect model generalization in a cross-species setting (see Table 5).

**TABLE 5.**
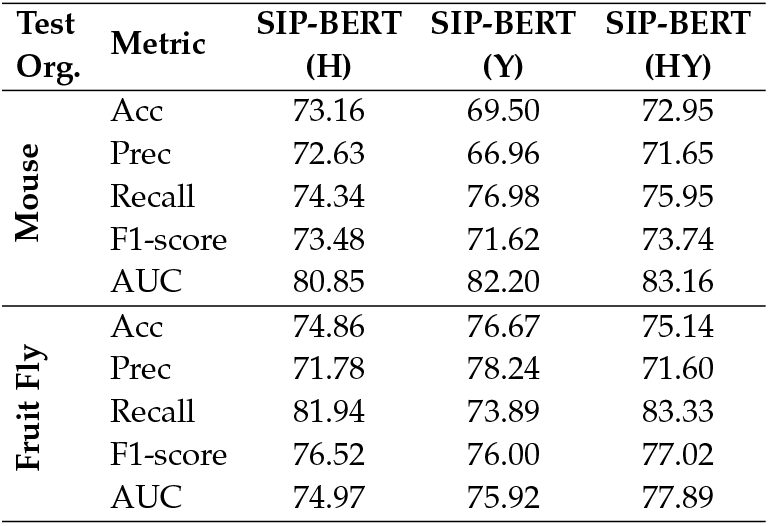
Cross-species evaluation of SIP-BERT models on balanced test sets from mouse and fruit fly.

### 3.4 Cross-Organism Performance Evaluation

A common limitation in many species represented in PPI databases is the scarcity of labeled self-interacting proteins (SIPs), which constrains the development of organism-specific predictive models. To address this, we investigated the cross-organism generalization capability of the proposed SIP-BERT models, SIP-BERT(H), SIP-BERT(Y), and SIP-BERT(HY), by evaluating their performance on two unseen species: mouse and fruit fly. These models were trained exclusively on human, yeast, or combined datasets and were not exposed to data from the test organisms during training.

The results, reported in Table 5, indicate that all models exhibit strong generalization across species boundaries. *SIP-BERT(HY)*, trained on both human and yeast data, achieved the highest performance (%) in fruit fly, with an AUC of 77.89 and an F1 score of 77.02. For the mouse, both *SIP-BERT(H)* and *SIP-BERT(HY)* yielded competitive results, attaining AUCs of 80.85 and 83.16, respectively. Importantly, the recall values remained consistently high across all settings, particularly for fruit fly, where they exceeded 81% for most models.

### 3.5 Performance Evaluation on Unbalanced Test Data

In our study, initial evaluations of model performance were carried out using balanced test sets to ensure fair bench-marking across methods. However, such balanced conditions rarely reflect the characteristics of real-world biological datasets, where non-self-interacting proteins (non-SIPs) vastly outnumber self-interacting proteins (SIPs) a trend consistently observed across both SIP-specific and general protein–protein interaction databases.

To evaluate the robustness and real-world applicability of our models under biologically realistic settings, we contructed five progressively unbalanced test sets with SIP-to-nonSIP ratios of 1:1, 1:2, 1:3, 1:4, and 1:5. Figure 3 illustrates radar plots summarizing five key evaluation metrics for three organism-specific SIP-BERT models (SIP-BERT(H), SIP-BERT(Y) and SIP-BERT(HY)) across these class imbalance conditions.

**Fig. 3.**
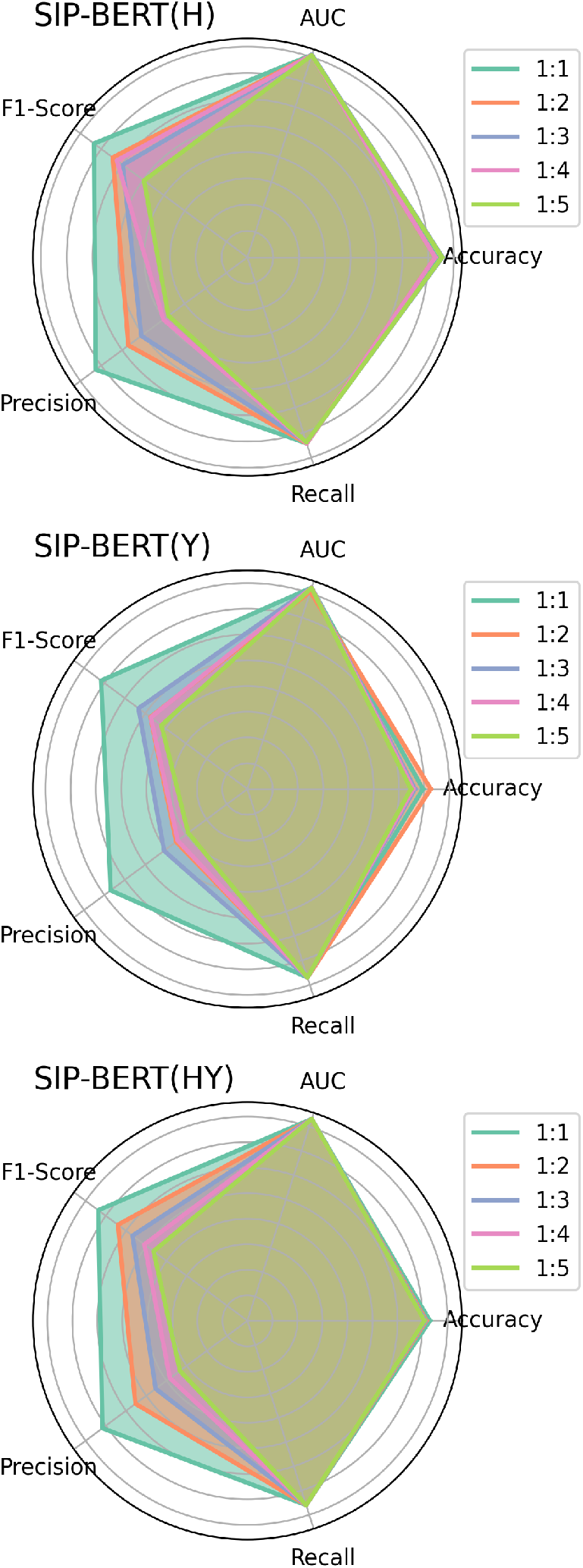
Vertically stacked radar charts illustrating the performance of SIP-BERT models across varying class imbalance ratios for SIP-BERT(H), SIP-BERT(Y) and SIP-BERT(HY).

The plots reveal a consistent trend: while Recall remains largely stable across increasing imbalance, both Precision and F1-score degrade notably, especially in the SIP-BERT(Y) model. This reflects the model’s tendency to preserve sensitivity (recall) at the cost of precision under skewed distributions. Despite this, the AUC and Accuracy metrics remain relatively stable, suggesting that SIP-BERT models maintain a strong discriminative capability even as class imbalance grows. These findings underscore the practical robustness of SIP-BERT models in diverse and imbalanced biological settings.

### 3.6 Structural Validation of Predicted False Positives Using PDB-Derived Contact Maps

We investigated a subset of false positive predictions, classified as self-interacting proteins (SIPs) by SIP-BERT(H) but lacking documented evidence in current PPI databases. To evaluate the structural plausibility of these predictions, we retrieved corresponding protein structures from the Protein Data Bank (PDB) [36] and generated *α − carbon*-based (*C*_*α*_ *− C*_*α*_) distance heatmaps to visualize inter-residue proximities. For three-dimensional structural assessment,we developed a `PyMOL` script [37] that highlights the entire protein structure in yellow while marking residues in close contact (5*≤* Å) in green. Figure 4 presents an example visualization of the structural assessment.

**Fig. 4.**
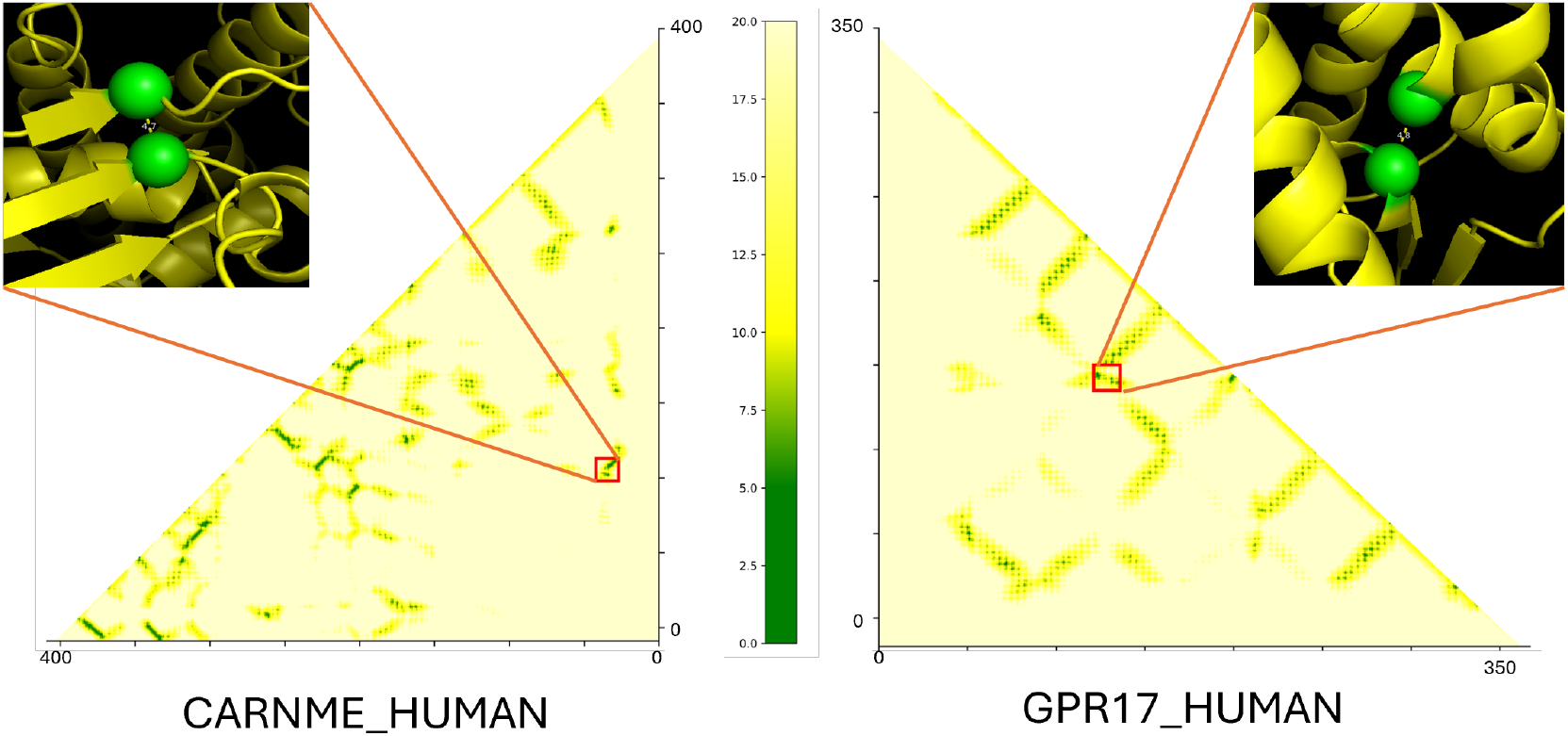
Residue-level *C*_*α*_–*C*_*α*_ distance heatmaps derived from experimentally resolved PDB structures for two proteins—*CARNME HUMAN* and *GPR17 HUMAN*—predicted as self-interacting by SIP-BERT(H), despite lacking annotations in current PPI databases. The x- and y-axes represent residue indices along the protein sequence, and the colour scale indicates inter-residue distances in angstroms (Å). Darker green regions correspond to residue pairs within 5 Å, highlighting potential intramolecular contacts suggestive of functional or structural self-interactions. Insets display zoomed-in 3D structural regions associated with high-contact zones, supporting the model‚s predictive ability to identify plausible, yet previously unannotated, self-interactions.

Notably, the heatmaps reveal substantial regions with alpha-carbon distances below 5 Å, indicating tight spatial proximity that is characteristic of potential self-interactions. These findings suggest that some of the false positive predictions may correspond to valid, yet unannotated, SIPs not currently captured by existing PPI databases.

## 4 Conclusion

In this study, we introduce SIP-BERT, an organism-independent deep language model designed for the prediction of protein self-interactions. SIP-BERT achieves superior performance compared to *state-of-the-art* methods, demonstrating accuracy improvements of 8% for S. cerevisiae (Table 2), 18% for human data, and 15% for a combined dataset of S. cerevisiae and human proteins. Notably, SIP-BERT comprises approximately 9 million trainable parameters, making it significantly more lightweight than ProteinBERT, which contains 16 million parameters.

SIP-BERT is particularly effective for cross-species prediction in scenarios where labeled data is scarce. When applied to underrepresented organisms such as *Mus musculus* (Mouse) and *Drosophila melanogaster* (FruitFly), the model achieves recall values exceeding 80%, highlighting its robust transferability across evolutionary lineages.

To further assess model predictions, we analyzed a subset of false-positives cases predicted as SIPs by SIP-BERT(H) but lacking supporting evidence in current PPI databases. Structural investigation using alpha-carbon distance heatmaps from the Protein Data Bank (PDB) revealed spatial proximity (5 *≤* Å) among residues, indicating potential self-interactions. These results suggest that SIP-BERT may uncover biologically plausible interactions not yet annotated in existing resources.

A key contribution of this work is the compilation and release of curated self-interaction datasets for multiple organisms, addressing the current scarcity of publicly available SIP data.

Future directions include validating novel predictions experimentally, extending SIP-BERT to additional species, and exploring more expressive architectures such as structure-informed transformers and contrastive learning-based multi-modal fusion. Together, these contributions advance the computational landscape for identifying self-interacting proteins and deepen our understanding of their functional roles across diverse biological systems.

## Acknowledgment

This work is partially supported by the CMATER research laboratory of the Department of Computer Science and Engineering, Jadavpur University, India. Subhadip Basu acknowledges the Department of Biotechnology grant (BT/PR16356/BID/7/596/2016) along with a Science and Engineering Research Board grant (SUR/2022/002903), Government of India.

## Author Declarations

The authors declare no conflict of interest. The dataset and source code are publicly available at https://github.com/CMATERJU-BIOINFO/SIP-BERT to promote open science and reproducibility. Tapas Chakraborty and Saikat Majumder contributed equally to this work as joint first authors. Tapas was responsible for the literature review and preparation of the initial manuscript draft, while Saikat implemented the code and conducted the experiments. Padmalochan Jena developed the datasets used in the study. The core research idea was jointly conceived by Subhadip Basu and Anup Kumar Halder, who also collaboratively formulated the scientific objectives and provided overall supervision throughout the research. All authors contributed to writing, reviewing, and approving the final version of the manuscript.

---

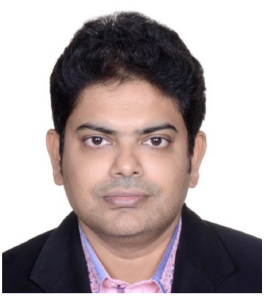

**Tapas Chakraborty** is pursuing his PhD under the supervision of Prof. Subhadip Basu from Jadavpur University, Kolkata, India. Tapas completed Bachelor of Engineering (in 2003) and Master of Engineering (in 2019) both in Computer Science and Engineering from Jadavpur University. He received a gold medal in board examination from Govt. of West Bengal for obtaining the highest marks in Mathematics. Tapas worked in the software industry for more than ten years. His research area includes Bio-informatics, Deep Learning, Audio and signal processing.

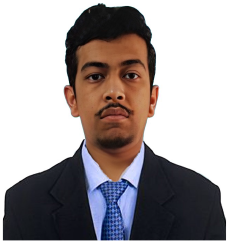

**Saikat Majumder** is currently working as a Junior Research Fellow (JRF) under the supervision of Prof. Subhadip Basu in the Department of Computer Science and Engineering at Jadavpur University, West Bengal, India. He completed his Master of Technology in Data Science at the Defense Institute of Technology, with expected completion in 2025. He completed his M.Sc. in Computer Science from the University of Delhi, Delhi, India, and previously earned his B.Sc. (Hons.) degree in Computer Science from Ramakrishna Mission Vidyamandira, affiliated with the University of Calcutta, West Bengal, India. His research interests include computational biology, bio-informatics, natural language processing, machine learning, and deep learning..

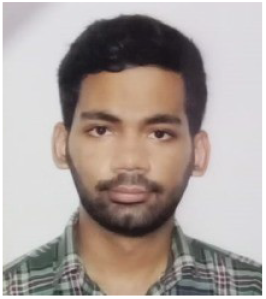

**Padmalochan Maiti** is currently pursuing his Bachelor of Engineering degree in Computer Science and Engineering from Jadavpur University, graduating in 2025. His areas of interest include machine learning, data science, and deep learning.

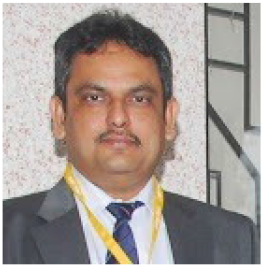

**Krishna Murthy** Dr. Krishna Murthy is currently Professor of Applied Mathematics at the Defence Institute of Advanced Technology (DIAT), Pune, India. He obtained his Ph.D. in Mathematics from the Indian Institute of Technology Kanpur in 2010 and subsequently held a Post-Doctoral Fellowship at École Centrale Paris,France, under the Erasmus Mundus–WillPower Fellowship. With over 11 years of teaching and research experience, his areas of interest include finite element analysis, computational fluid dynamics, mathematical modelling, numerical methods for PDEs, cryptography, machine learning, and image processing. Dr. Murthy has contributed to various national and international journals and conferences, serves as Editor-in-Chief of the International Journal of Thermal Energy and Applications, and is on editorial boards of other professional journals. He has active collaborations with DRDO labs, academic institutions, and industries both in India and abroad.

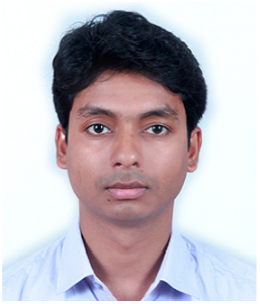

**Anup Kumar Halder** (M’24) is an Assistant Professor in the Department of Computer Science and Engineering at the Indian Institute of Information Technology, Raichur, India. He previously held the position of Assistant Professor of Research (Postdoctoral Fellow) at the Laboratory of Functional and Structural Genomics, Centre of New Technologies (CeNT), University of Warsaw, and at the Faculty of Mathematics and Information Science, Warsaw University of Technology, Poland. He earned his Master’s and Ph.D. degrees in Computer Science and Engineering from Jadavpur University in 2014 and 2021, respectively. A recipient of the prestigious Visvesvaraya PhD Fellowship from the Government of India, his research focusses on deep learning and machine learning applications for biological data analysis, with expertise in proteomics, 3D genomics, biological networks, and pattern recognition.

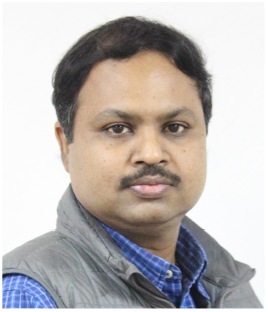

**Subhadip Basu** (SM’12) is a Professor in the Department of Computer Science and Engineering at Jadavpur University, India. He received Ph.D. in Engineering from Jadavpur University in 2006 and has more than 200 research publications in notable Journals and Conferences in the areas of bioinformatics, image analysis, pattern recognition, etc. Dr. Basu is the recipient of DAAD fellowship from Germany, BOYSCAST fellowship, and UGC Research Award from the Govt. of India, HIVIP fellowship from Hitachi,Japan, EMMA and cLink fellowships from the European Union.

